# Cadmium tolerance is associated with tissue-specific plasticity of metallothionein gene expression in *Gammarus fossarum* field populations

**DOI:** 10.1101/2025.01.09.632096

**Authors:** Auréline Lalouette, Arnaud Chaumot, Louveline Lepeule, Karen Gaget, Nicolas Delorme, Laura Garnero, Federica Calevro, Davide Degli Esposti

## Abstract

The metallothionein gene family codes for proteins involved in metal homeostasis and acute detoxification of non-essential toxic metal ions across the tree of life. We have previously documented increased cadmium (Cd) tolerance in field populations of the crustacean *Gammarus fossarum* exposed to chronic metallic contamination of geochemical origin. This tolerance is lost during maintenance of organisms in the laboratory, and is transmitted to offspring via parental effects. This study investigated whether the expression of the Cd-responsive metallothionein gene *mt1* could be related to Cd-tolerance plasticity in *G. fossarum*. In eleven populations with different chronic Cd exposure history, we simultaneously assessed Cd-tolerance (mortality tests) and *G. fossarum mt1* expression levels by RT-qPCR in the gills and caeca of adult males and in neonates. *mt1* expression levels in the two organs were correlated to Cd-tolerance in field organisms and a loss of tolerance was observed in parallel with a decreased expression of *mt1* in the caeca after maintenance in uncontaminated water. We also recorded a greater inducibility of *mt1* expression in offspring of tolerant populations in the laboratory when re-exposed to Cd along with the bi-parental transmission of Cd-tolerance. These results suggest that the control of *mt1* expression is involved in the plasticity of Cd-tolerance in gammarid populations with different histories of Cd exposure.

**Highlights:**

- *mt1* gene expression in the gills and in the hepatopancreatic caeca is associated with Cd tolerance in *Gammarus fossarum* populations.
- Plasticity of *G. fossarum mt1* gene expression is associated to loss of Cd tolerance.
- Parental effect in Cd tolerance is supported by transgenerational control of *mt1* expression, underlid by an increased induction of *mt1* during Cd exposure.

## 1. Introduction

In aquatic ecosystems, heavy metal contamination is a major concern as they are continuously released into the environment either through anthropogenic or natural processes (Garrett, 2000). In response to variable metal contamination in the environment, organisms can develop increased tolerance to metals (*i*.*e*. changes in sensitivity), notably through the development of transgenerational acclimation mechanisms (Morgan 2007; Amiard-Triquet 2019). Tolerance to metal exposure may involve different metal regulation mechanisms that notably detoxify and eliminate metallic elements that are incorporated into the organism (Amiard-Triquet, 2019). Among these mechanisms, the expression of members of the metallothionein (*mt*) gene superfamily has been documented to confer tolerance to metal exposure (Amiard et al., 2006; Dallinger and Höckner, 2013). MTs are cysteine-rich, low molecular weight proteins involved in essential metal homeostasis, non-essential metal detoxification and protection against oxidative stress (Amiard et al., 2006; Mao et al., 2012). These proteins are highly conserved across the animal kingdom and are considered to be the main defence against toxic metals in invertebrates (Ziller and Fraissinet-Tachet, 2018; Duarte et al., 2019). On the other hand, the involvement of MTs in metal tolerance has been suggested in many terrestrial and aquatic species (as shown in reviews by Amiard et al., 2006; Morgan et al., 2007; Janssens et al., 2009; Dallinger and Höckner, 2013). A higher basal level and/or a stronger induction capacity of MTs have been observed in tolerant populations than in naïve populations of the same species. For example, higher levels of MTs were found in cadmium (Cd) tolerant populations of *Chironomius riparius* compared with susceptible populations of the same species after field sampling (Pedrosa et al., 2017). In contrast, in *Orchesella cincta*, similar levels were found between Cd tolerant and sensitive populations in the field, but with a greater capacity to induce MT in tolerant populations when they are re-exposed to the metal of interest (Sterenborg and Roelofs, 2003). Furthermore, the involvement of MT in the transmission of tolerance to offspring has been little studied, particularly in invertebrates. Nevertheless, the potential role of MT in the transfer of metal tolerance to offspring has been suggested in some studies conducted in daphnia (Tsui and Wang, 2005; Guan and Wang, 2006), bivalves (Weng and Wang, 2014) and fish (Wu et al., 2008, 2012).

Species of the genus *Gammarus* have been studied in the context of the acquisition of metal-induced tolerance and of the long-term effects of metal exposure (*e*.*g*. Stuhlbacher and Maltby, 1992; Khan et al., 2011; Vigneron et al., 2015). Recently, we have shown that Cd tolerance, can be observed in several populations of *G. fossarum* living in rivers naturally contaminated by geochemical sources of Cd (Lalouette et al., 2023, 2024). In these populations, tolerance appears to be linked to non-genetic acclimation, and more specifically to phenotypic plasticity, since this tolerance can be temporarily lost in populations and by individuals when they are isolated in Cd-free water (Lalouette et al., 2023). Furthermore, Cd tolerance is transmissible to offspring through parental effects, seems to depend on long-term parental exposure, and can be transmitted by both mother and father (Lalouette et al., 2023). However, the molecular mechanisms mediating these observations are not currently known for this species. Otherwise, a recent work on this species highlighted the toxicokinetic of Cd ions in various organs (gills, caeca, intestine, cephalon and other tissue) (Gestin et al., 2022). Specifically, Gestin et al. (2022) showed that intestines are involved in the uptake, distribution and elimination of Cd, that caeca play a central in accumulation and elimination function, while gills are hyper-accumulative organs with an important role in storage of Cd. Taking advantage of the results on the toxicokinetic of Cd and to the recent identification of a Cd-responsive metallothionein coding transcript (*mt1*) in the *G. fossarum* transcriptome (Degli Esposti et al., 2024), this study aimed to: (i) determine the association between *G. fossarum mt1* expression and Cd tolerance at different life stages and sub-individual (organ) and (ii) assess parental transmission of *G. fossarum mt1* expression and its effects on Cd-tolerance in the offspring.

## 2. Material and Methods

### 2.2. Selection of the eleven *G. fossarum* populations

Based on our previous works (Lalouette et al., 2024) we considered eleven populations of *G. fossarum* inhabiting upstream areas of headwater streams in natural or semi-natural context and presenting a contrast in exposure to Cd-contamination and Cd-tolerance. More precisely, we selected three populations living in rivers naturally contaminated by Cd and presenting tolerance to this metal (Ardillats, Marchampt and Vernay populations), three populations living in rivers presenting intermediate Cd-contamination and tolerance levels (Strenbach, Rauental and Morcille populations) and five populations living in uncontaminated rivers and presenting no tolerance to Cd (Seran, Doulonne, Katlen, Reigne and Vancelle populations). The results of the distribution of individual survival times obtained during acute Cd sensitivity tests on adult male gammarids freshly collected in the field during the sampling campaign of the present study are already supplied in Lalouette et al. (2024). Each population has been assigned a letter as it follows: Ardillats (A), Marchampt (B), Vernay (C), Strenbach (D), Rauenthal (E), Morcille (F), Séran (G), Doulonne (H), Katlen (I), Reigne (J) and Vancelle (K). Futhermore, an active biomonitoring approach (caging and bioaccumulation measurement in transplanted reference organisms) were used to assess populations’ field exposure to Cd as described in previous studies (Vigneron et al., 2015; Ciliberti et al., 2017; Alric et al., 2019). The average level of Cd contamination over the last ten years (already published in Lalouette et al. (2024)) and the location details of the eleven study sites are presented in Supplementary Table 1.

### 2.2. Sampling and maintenance of gammarids in the lab

Two sampling campaigns were realised for this study: one in June 2022 (eleven populations collected) and another in October 2022 (five populations collected). For each campaign, gammarids were collected by kick sampling and quickly transported to the laboratory in plastic buckets containing ambient freshwater. They were maintained in the water of their own site at 12°C and they were fed *ad libitum* with conditioned alder leaves (*Alnus glutinosa*) until used for the experiments (max 24h). The day after field sampling and for each population, 6 males per population were dissected and the gills and the caeca were individually collected for each organism as described in Gestin et al. (2021) to measure *mt1* gene expression (see section 2.6). Organs were flash-frozen in liquid nitrogen and kept at - 80°C until RNA extraction.

Gammarids were maintained in the laboratory in plastic aquariums (2 L or 11 L) placed in water baths thermos-regulated at 16°C and with a controlled photoperiod (16/8 night/day). Gammarids were fed *ad libitum* with conditioned alder leaves (*Alnus glutinosa*) and a weekly supply of Tubifex larvae.

### 2.3. Deacclimatation of populations in Cd-free water in the laboratory

Gammarid organisms from populations A, B, C, G and H were collected in October 2022 and were kept in separate 11 L plastic aquaria (around 1000 adult individuals). After 2 months, male adults from each population were collected for Cd sensitivity tests (see section 2.5) and to measure *mt1* gene expression in gills and caeca (see section 2.6) as previously described. In parallel, we sorted 100 gammarids in mating-pairs with females in the last stage of their reproductive cycle (stage D2 with hatched neonates in brood pouch) from the five populations collected in December 2022 (A, B, C, G, H). After one reproduction cycle (approximatively 25 days), neonates from each population were collected for Cd sensitivity tests (n = 24; see section 2.5) and to measure *mt1* gene expression (n = 6 pool of 10 neonates; see section 2.6). In this same batch of neonates, we studied MT inducibility by measuring *mt1* gene expression after re-exposure at sublethal levels to Cd. For this purpose and for each population, 120 neonates were collected and isolated by pool of 10 neonates in falcons containing a mixture of Evian water and demineralised water (50/50, conductivity approximately 300 μs/cm) with or without CdCl_2_ (equivalent to 3 μg/L of dissolved Cd^2+^). For each condition, 6 falcons were considered and exposure was conducted over 3 days. At the end of the 3 days, neonates from both conditions were collected, placed in 2 mL eppendorf tubes and frozen at -80°C.

### 2.4. Cross-breeding of tolerant and sensitive populations

In order to assess parental effects on *mt1* expression in the offspring, we carried out cross-breeding between genitors from the population A (tolerant) and population G (sensitive) collected during the campaign of June 2022 as described in Lalouette et al. (2023). Briefly, after sampled in the field, gammarids in mating-pairs were sorted visually with females in stage D2, separated and re-mated between tolerant and sensitive population, resulting in four conditions: AA, GG, AG, GA, where the first letters correspond to the initial of the population name of the male and the second to the initial of the population name of the female. For each condition, two 2-L plastic aquariums containing 50 mating-pairs of gammarids were considered. At the end of the moulting-cycle (approximatively 25 days), neonates from the different conditions were collected for Cd sensitivity tests and to measure the expression of *mt1* gene expression. Moreover, the inducibility of *mt1* was studied following re-exposure to Cd (3 µg/L) during 3 days using the same protocol as described previously (see section 2.3).

### 2.5. Acute toxicity tests for Cd tolerance assessment

Tolerance to Cd was assessed in adult males after 2 months in the laboratory. Similarly, tolerance was also measured in the cohort of neonates born in the laboratory from field populations or from crosses between populations. For that, we performed acute toxicity tests as described in Lalouette et al. (2023, 2024). For each condition of adult gammarids (source population, laboratory maintenance duration), three replicates of 15 males were exposed to a nominal concentration of 80 µg/L of Cd in 500 mL plastic beakers. For progeny tolerance assessment, 24 neonates to each condition were collected one day after their release from maternal marsupium. The acute toxicity tests were then carried out by individually monitoring the survival of neonates exposed to a nominal Cd concentration of 20 µg/L in 50 mL falcons®. As previously described (Lalouette et al., 2023, 2024), the different Cd solution was prepared from stock solutions of Cd at 0.1 g/L. Then, beakers and falcons® were kept in a thermo-regulated water bath at 12°C and mortality was monitoring daily during exposure tests.

### 2.6. Gene expression analysis

Gene expression of *mt1* was measured by RT-qPCR (Degli Esposti et al. 2024). Here it follows a description of the different steps.

#### Selection of the studied organs

We focused on *mt1* gene expression in gills and caeca for adults’ males gammarids. Indeed, a preliminary experiment was conducted in the gills, caeca, ovaries and testes of adult individuals from the tolerant (Ardillats) and the sensitive (Séran) population. The preliminary results showed differences between *mt1* gene expression in the caeca and gills, as well as in the ovaries, but not in the testes (Supplementary Figure 1). Since this study focuses only on male (to avoid confounding factors linked to the reproductive cycle of females), we chose to focus on the caeca and the gills, two organs involved in in Cd toxicokinetics in *G. fossarum*.

#### Sample

For adult males, we considered 6 biological replicates per organ and population. For neonates, *mt1* gene expression was measured in pools of 10 neonates. For each measurement, 6 pools of 10 neonates were considered. Then, the same protocol was used for adult and neonates.

#### Total RNA extraction and cDNA synthesis

The RNeasy fibrous tissues kit (Qiagen, Hilden, Germany) was used to extracted RNA from each organ or pool of neonates and quality and quantity were assessed by spectrophotometry using NanodropOne (Thermofisher, Waltham, MA, USA). RNAs (about 100 ng per sample) were retro-transcribed using the high-capacity cDNA reverse transcription kit (Applied Biosystems, Waltham, MA, USA).

#### Quantitative real-time polymerase chain reaction (qPCR)

For target genes, specific primers for *mt1* and *elongation factor (ef)* genes were used (Supplementary Table 2). Following the reverse transcription, cDNAs was diluted (1:5) and amplified using the OneGreen sybr green mix (Ozyme, Montigny-le-Bretonneux, France) in a Bio-Rad CFX96 qPCR platform (Hercules, CA, USA) using the following conditions: 95°C for 3 min followed by 40 cycles at 95°C for 10 s, 61°C for 30 s and 72°C for 30 s. Three technical replicates were performed for each sample. MTs were quantified using the relative quantification method (ΔCt) using the elongation factor (EF) expression to normalize gene expression across samples, as in Degli Esposti et al. (2024). The reference gene *ef* is a stable gene on *G. fossarum* (Gouveia et al., 2018). In order to obtain a better visualization of the results, these were transformed using the square root function (√ΔCt).

### 2.7. Statistical analysis

The data were processed with the R software version 4.4.0 (R Core Team, 2024). The data obtained during acute toxicity tests (sensitivity to Cd) were analysed by Kaplan-Meier type survival analyses (log-rank test) using the “survival” R package. For *G. fossarum mt1* expression, the results obtained were analysed using the Pairwise Wilcoxon rank sum test (abbreviation used: PW) with correction for multiple testing by the Benjamini-Hochberg method. All the statistical analyses were performed with a statistical significance level of 5%.

## 3. Results

### 3.1. Basal level of *G. fossarum mt1* gene expression in adults sampled in field

The expression of *mt1* was measured in the gills and the caeca of adult males after field sampling in tolerant, intermediate and sensitive populations (Figure 1, Supplementary Figure 2). The results shown that *mt1* expression is 8-fold higher in the gills of organisms from contaminated populations than in uncontaminated (PW test: adjusted p-value = 3.10^−9^) and 5-fold higher than those from intermediate populations (PW test: adjusted p-value = 0.03) (Figure 1, Supplementary Figure 2). Similarly, the intermediate populations showed higher levels of *mt1* expression than the uncontaminated populations (PW test: adjusted p-value = 9.10^−9^). In the caeca, *mt1* expression levels (Figure 1) of organisms from contaminated populations were 2-fold higher than those from uncontaminated populations (PW test: adjusted p-value = 8.10^−5^). Interestingly, the results reveal a positive correlation between the median survival time (LT50) of each population and the median expression of *mt1* in the gills (Spearman-correlation, ρ = 0.8; p-value = 10^−3^; Figure 2) and caeca (Spearman-correlation, ρ = 0.7; p-value = 10^−2^; Figure 2) of adult males.

**Figure 1:**
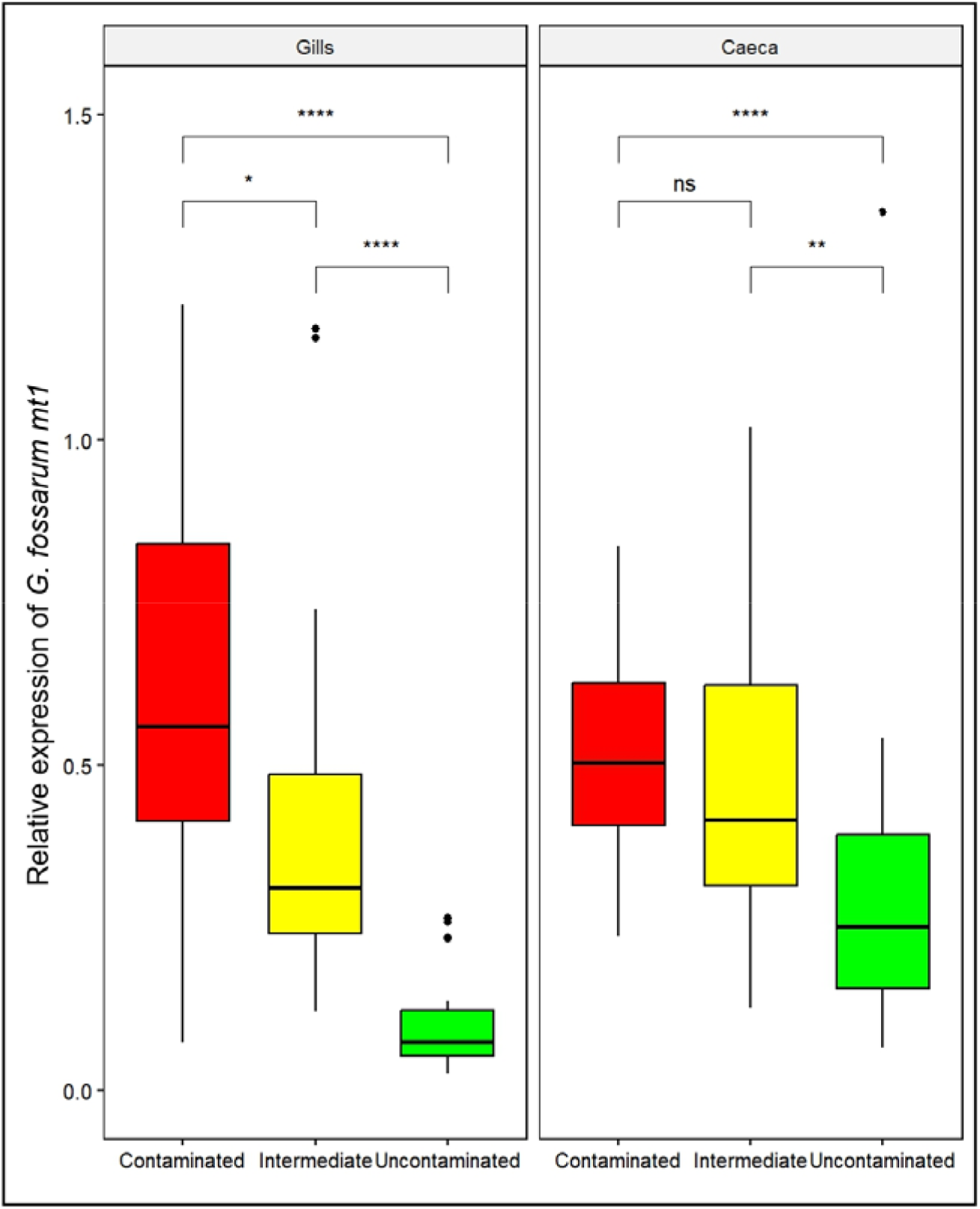
*G. fossarum mt1* gene expression measured by RT-qPCR in the gills and caeca of adult males after sampled in the field in contaminated (pooled data from populations A, B and C), intermediate (pooled data from populations D, E and F) and uncontaminated (pooled data from populations G, H, I, J and K) populations. Gene expression was normalize using the elongation factor (EF) C_t_ value. Data detailed by population are presented in Supplementary Figure 1. Red, yellow and green colors correspond respectively to Cd contaminated, intermediate and uncontaminated status of populations. The symbols indicate the statistical significance (p-value) as followed: (ns) > 0.05; (*) ≤ 0.05; (**) ≤ 0.01; (***) ≤ 0.001; (****) ≤ 0.0001.

**Figure 2:**
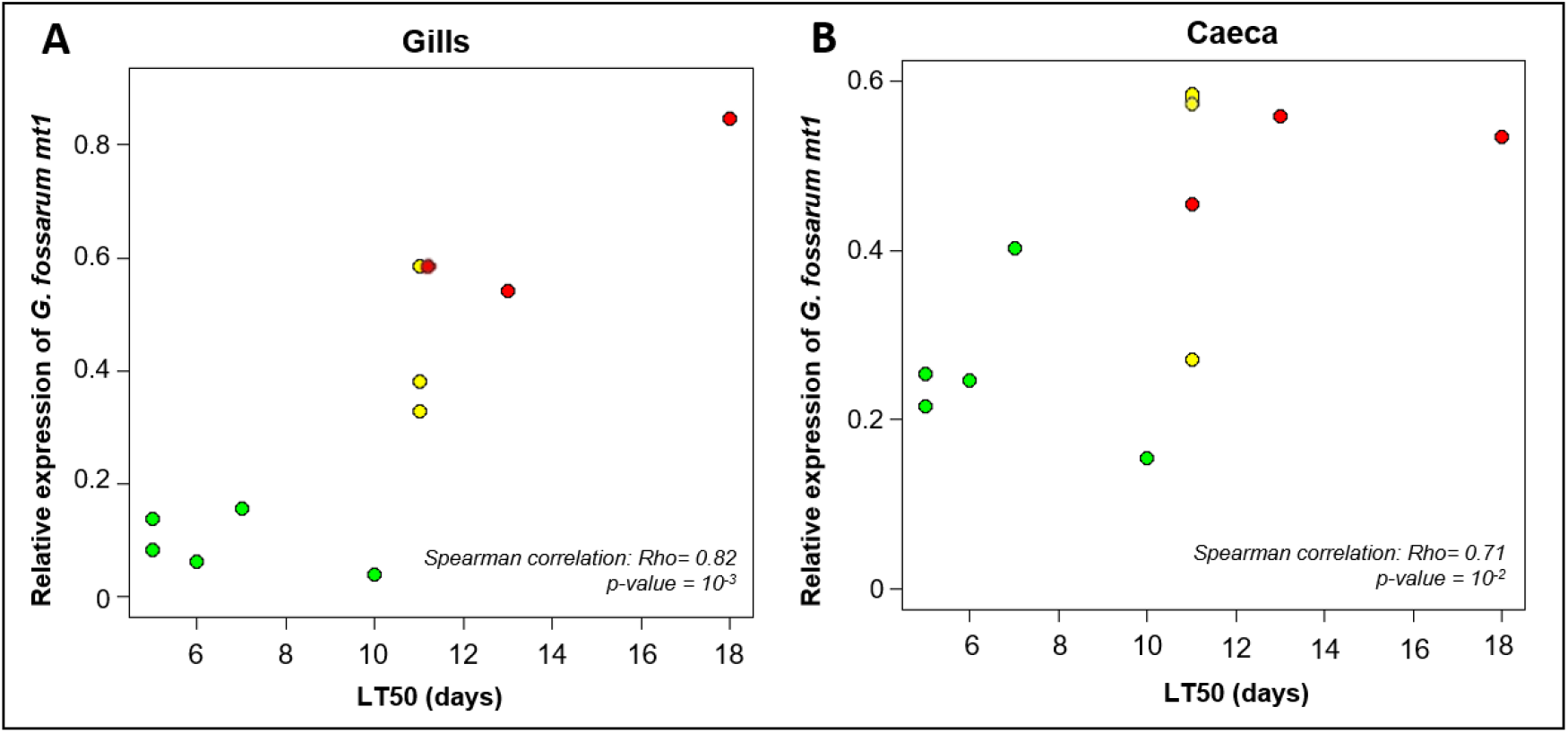
Correlation plots between the average relative expression of *G. fossarum mt1* in gills (**A**) and caeca (**B**) with the median (LT50) of adults survival time (days) of the 11 populations. Red, yellow and green colors correspond to Cd contaminated, intermediate and uncontaminated status of populations, respectively (see Supplementary Table 1).

### 3.2. Plasticity of *G. fossarum mt1* gene expression is associated to loss of Cd tolerance

After two months of maintenance in Cd-free water (T2), we observed a decrease in Cd tolerance compared to those observed just after field sampling (T0) in population A (log-rank test: p-value = 6.10^−3^) and population C (log-rank test: p-value = 8.10^−8^) (Supplementary Figure 3). In particular, the decrease in survival time distribution was driven by a loss of the highest levels of individual tolerance, together with a decrease of the lowest levels down to the range of survival times observed in uncontaminated populations (Supplementary Table 3). For example, only 16% and 14% of the A-T0 and C-T0 individuals respectively, had a mean survival time inferior or equal to 6 days compared to 40% of A-T2 and 44% of C-T2 (Supplementary Table 3). For population B, no tolerance to Cd was observed in gammarids after field sampling at the date tested. Thus, no difference in Cd sensitivity was observed between B-T0 and B-T2.

Gene expression levels of *mt1* in the gills and caeca of adult males before and after the maintenance in Cd-free water were also measured (Figure 3). There was a significant decrease in *mt1* expression in the caeca of organisms from populations A and B (Pairwise-Wilcoxon (PW) test: adjusted p-values < 0.03). The same trend was observed, but not significantly, for organisms from population C (PW test: adjusted p-value = 0.58). In contrast, there was no modulation of *mt1* expression in the gills of organisms from these same populations after 2 months in clean water (PW test: adjusted p-values > 0.4). For uncontaminated populations, no difference in *mt1* expression in the caeca or gills was observed over time (PW test: adjusted p-value = 0.26 and p-value= 0.77, respectively).

**Figure 3:**
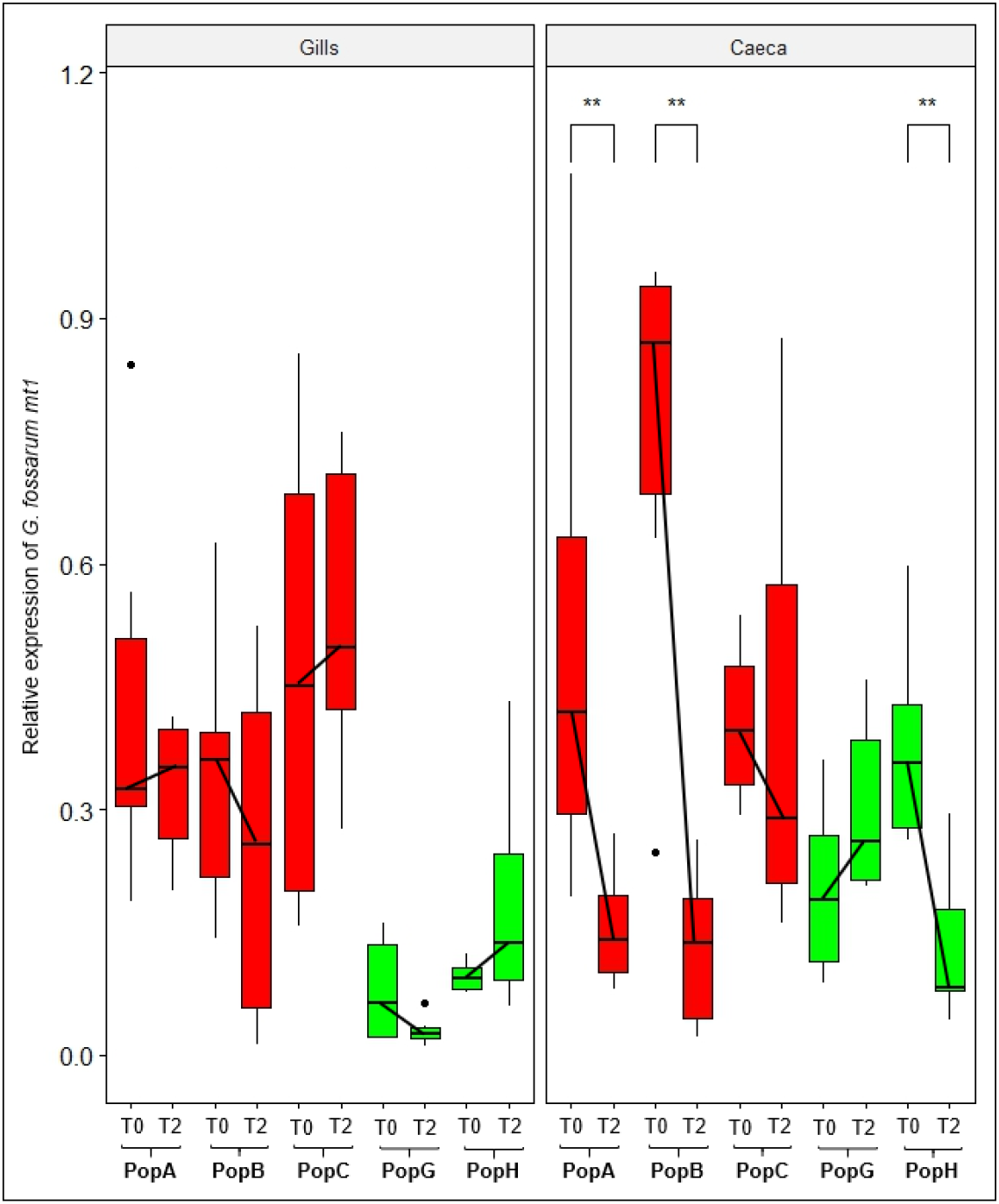
*G. fossarum mt1* gene expression measured by RT-qPCR in the gills and caeca of adult males after sampled in the field (T0) and after maintenance in Cd-free water during 2 months in the lab (T2) in five field populations (A, B, C, G, H). Gene expression was normalize using the elongation factor (EF) C_t_ value. Red and green colors correspond respectively to Cd contaminated and uncontaminated status of populations in their habitat of origin. The symbols indicate the statistical significance (p-value) as followed: (ns) > 0.05; (*) ≤ 0.05; (**) ≤ 0.01.

### 3.3. *G. fossarum mt1* gene expression and Cd tolerance in the offspring

In order to investigate the relationship between *mt1* gene expression and Cd tolerance in the offspring, we focused on the offspring produced in the lab by genitors exposed on the field and kept in Cd-free water. The results of acute Cd sensitivity tests showed that neonates from the contaminated populations were more tolerant compared to those from the uncontaminated populations (log-rank test: p-value = 10^−12^) (Figure 4A, Supplementary Figure 4). However, no difference in *mt1* expression was observed between the tolerant and sensitive populations of neonates before the acute exposure (PW test: adjusted p-value = 0.72) (Supplementary Figure 5). Thus, we tested the capacity of neonates to induce MT by exposing them to 3 µg/L Cd for 3 days, followed by measurement of *mt1* expression. Results showed that *mt1* inducibility is higher in tolerant neonates (Figure 4B, Supplementary Figure 5). Neonates from tolerant parents were able, on average, to induce *mt1* expression four times (Fold-change = 8) more strongly than uncontaminated populations (Fold-change = 2).

**Figure 4:**
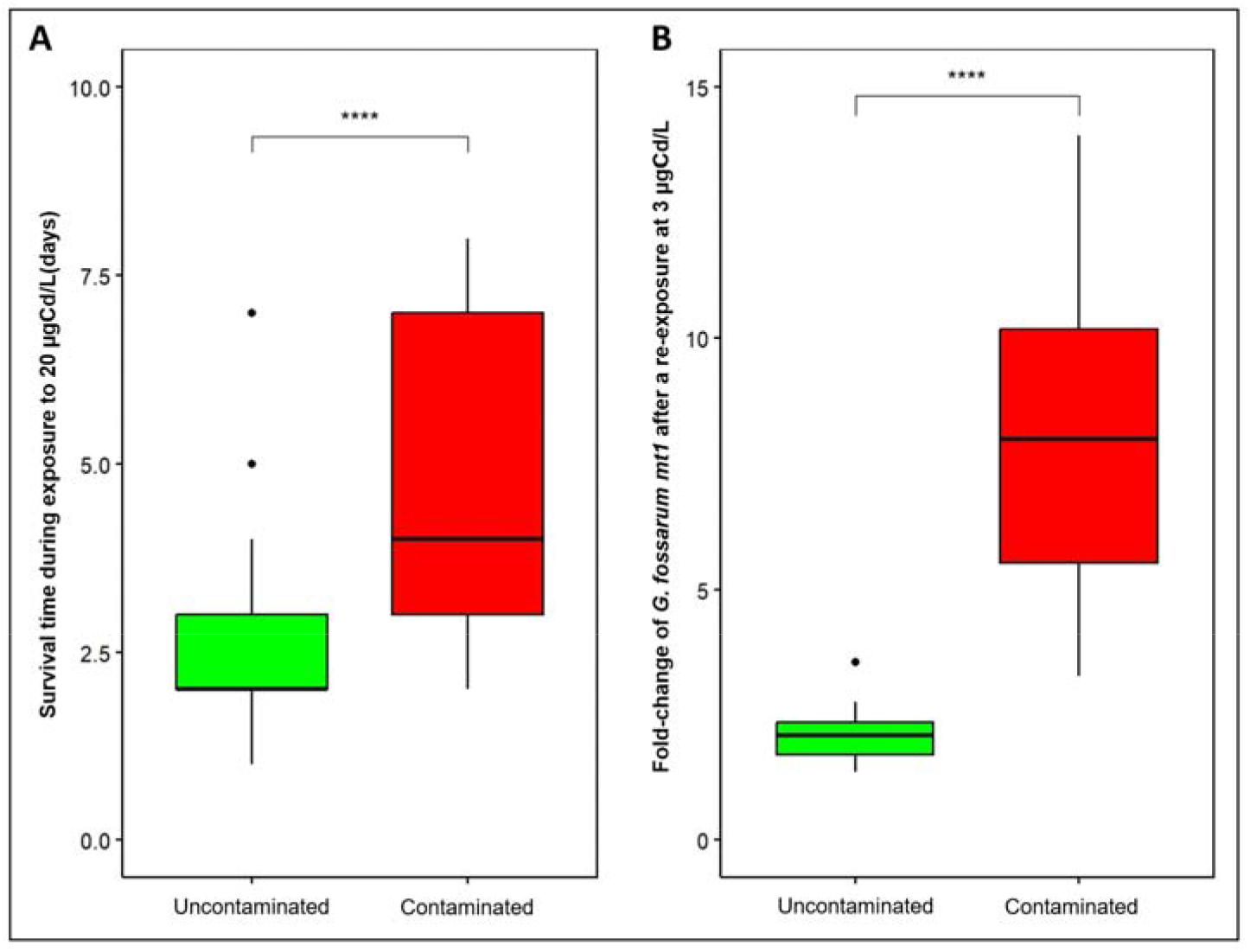
Tolerance and *mt1* expression levels in neonates born in the lab in Cd free water after one reproduction cycle (approximatively 25 days) from three contaminated populations and three uncontaminated populations. (**A**) Survival time of offspring exposed to Cd (20 µg/L) individually one day after their release from maternal marsupium and (**B**) Fold change of *G. fossarum mt1* gene expression measured by RT-qPCR in neonates after a re-exposure at 3 µgCd/L during 3 days compared to non-exposed controls. Data detailed by population are presented in Supplementary Figure 3 and 4. Red and green colors correspond respectively to Cd contaminated and uncontaminated status of populations in their habitat of origin. The symbols indicate the statistical significance (p-value): (****) ≤ 0.0001.

**Figure 5:**
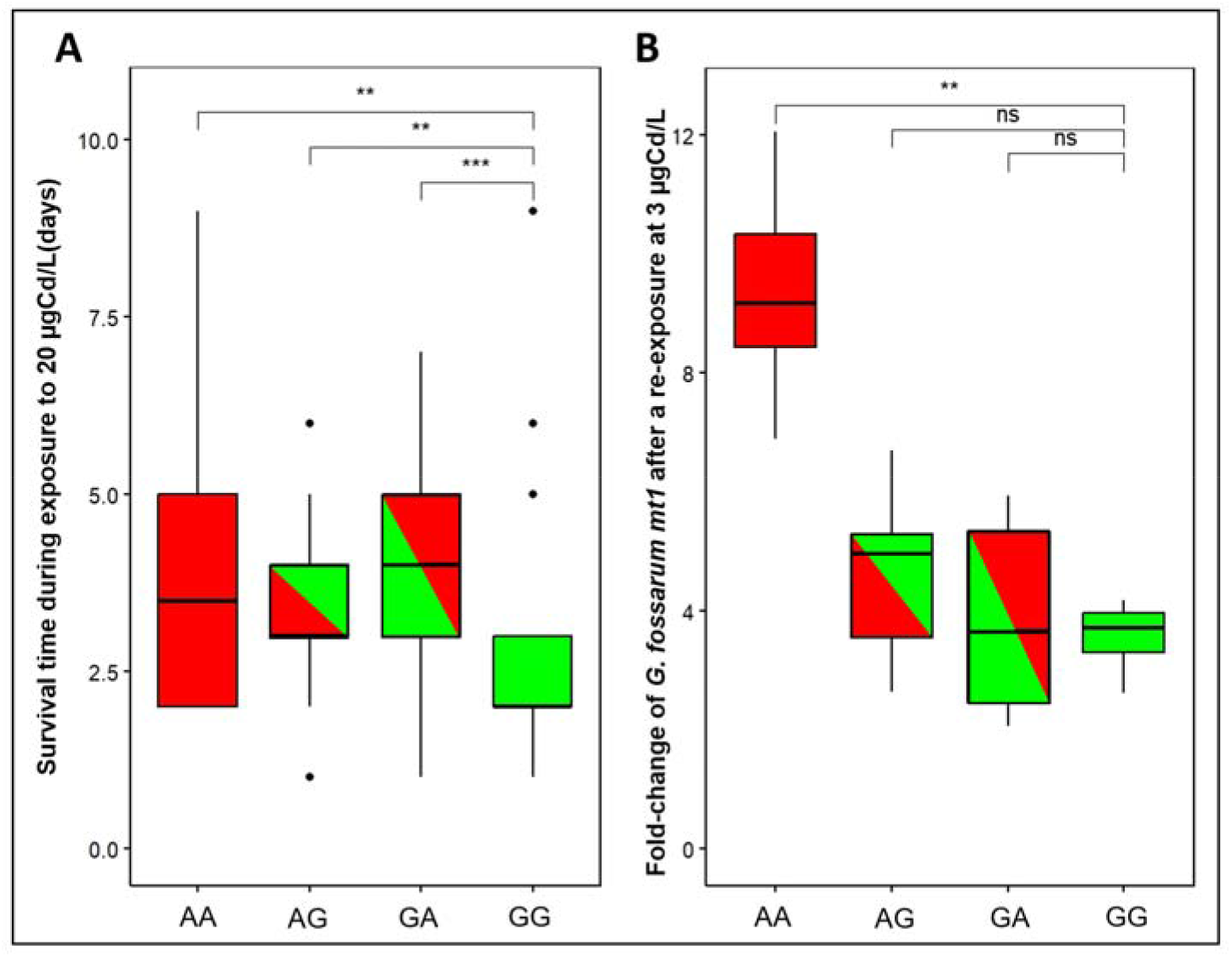
Cross-breeding between sensitive and tolerant populations. The figure presents (**A**) the Cd tolerance (distribution of individual survival times) and (**B**) the fold-change of *G. fossarum mt1* gene expression measured by RT-qPCR after re-exposure at 3 µgCd/L during 3 days in neonates produced by Cd tolerant population (population A), uncontaminated population (population G) and from population cross breeding. For the legend of the different conditions, the first capital letter corresponds to the first letter of the male’s population name and the second capital letter correspond to the first letter of the female’s population name (notation ♂**/**♀: *e*.*g*, the condition AG corresponds to the crossing between a male from A population and a female from G population). The symbols indicate the statistical significance (p-value) as followed: (ns) > 0.05; (*) ≤ 0.05; (**) ≤ 0.01; (***) ≤ 0.001.

Finally, crossbreeding experiments between parents issued from populations A and G showed that neonates born from a tolerant father (condition AG) and from a tolerant mother (condition GA) present a distribution of survival times intermediate between the distributions from the two parental population conditions (AA and GG) (Figure 5A). Indeed, the distribution of survival times of GA neonates is not significantly different from AA condition (log-rank test: p-value = 0.3) while the AG condition differs significantly to GG condition (log-rank test: p-values = 6.10^−3^). The expression levels of *mt1* were measured in control condition and after a challenge at 3 µgCd/L for 3 days (Figure 5B, Supplementary Figure 6). The results show that *mt1* inducibility was higher in AA and AG neonates (Fold-change = 9.2 and 5, respectively) under Cd conditions than in GG neonates (Fold-change = 3.7) (Figure 5B). In particularly, we observed an increase in individual variability in neonates AG and GA, with respectively, 67% and 50% of individuals presenting a fold-change greater than 4, against 100% for AA condition and 17% for GG condition (Supplementary Table 4).

## 4. Discussion

The ability of populations to maintain themselves in metal-contaminated environments depends on the effectiveness of mechanisms that protect the organisms from metal uptake and/or contribute to metal detoxification (Stohs, 1995). One of these mechanisms involve the expression of genes belonging to the metallothionein (*mt*) superfamily. Several studies have suggested that metal detoxification by metallothioneins is associated with increased tolerance to metals (Morgan et al., 2007). The implication of MTs in response to metal exposure and/or in metal tolerance has also been documented in various arthropod species such as amphipods (Stuhlbacher and Maltby, 1992; Martinez et al., 1996; Correia et al., 2002; Dong et al., 2020), daphnids (Tsui and Wang, 2005; Guan and Wang, 2006; Li et al., 2016) or diptera (Mireji et al., 2010; Rosabal et al., 2012). In two species of arthropods, *Chironomius riparius* and *Orchesella cincta*, tolerant individuals from metal-contaminated sites show increased expression of *mt* gene compared with a reference population (Pedrosa et al., 2017; Timmermans et al., 2005; Roelofs et al., 2009). The up-modulation of MTs expression is considered to be part of antioxidant processes by binding free metals (like Cd or Pb) and thus reducing their pro-oxidant effects (Amiard et al., 2006; Mao et al., 2012). Similarly, in tolerant populations, it has been suggested that increasing baseline levels of MTs would help to maintain homeostasis in these populations (Atrian and Capdevila, 2013). However, establishing this link is challenging in field populations since the majority of these studies in the literature compare only one tolerant population *versus* one sensitive population (Dong et al., 2020; Chiodi-Boudet et al., 2019; Sterenborg et al., 2003). In fact, such experimental designs are subject to possible false positive conclusions in relation to the potential influence of confounding biotic or abiotic factors unrelated to contaminant exposure. Thus, we investigated the link between the expression of *G. fossarum mt1* described as being specifically induced during short-term Cd exposure in *G. fossarum* (Degli Esposti et al., 2024) and the plasticity of Cd tolerance observed in chronically exposed natural populations of this species (Lalouette et al., 2023). Firstly, we measured *mt1* expression in Cd-tolerant and Cd-sensitive natural populations after field sampling in two organs, the caeca and the gills, known to be involved in Cd toxicokinetics in *G. fossarum* (Gestin et al., 2022). Our results showed a correlation between *mt1* expression and Cd tolerance, with higher levels of *mt1* expression in the gills and caeca of tolerant populations. The strength of our study is to consider eleven field populations with similar living habitats except for Cd-exposure, thus limiting the impact of other environmental confounding factors (Lalouette et al., 2024).

Our results also showed that in gammarids exposed to Cd in natural environment, *mt1* is differentially expressed in populations with different histories of metal exposure. More specifically, we showed that the average expression of *mt1* is approximately 4 and 6 times higher in the gills of intermediate and tolerant populations, respectively, compared with sensitive populations. Similarly, in the caeca, the average expression was 2 times higher in the intermediate and tolerant populations. Similar results were found in the gills of the mussel *Pyganodon grandis*, where MT concentrations (measured as protein fraction) increased along the gradient of metal contamination at the different study sites (Bonneris et al., 2005). However, a correlation between tolerance and MT expression is not systematically found. For example, Chiodi Boudet et al. (2019) showed different detoxification strategies in *Palaemon argentinus*. When this species is exposed to metals, the main detoxification pathway in tolerant populations is Cd complexation in granules (Cd-MRG), whereas sensitive populations mainly detoxify Cd by binding to MTs. Similarly, Vijver et al. (2004) suggest that granules are responsible for tolerance during long-term exposure to Cd, whereas MTs are involved mainly during short-term exposure.

Afterwards we investigated whether Cd tolerance plasticity was mirrored by *mt1* expression changes. For that, three tolerant populations (population A, B, C) and two sensitive populations (population G, H) were maintained in Cd free water in the lab for 2 months. When individuals from Cd-tolerant populations were no longer exposed to this metal for 2 months, there was a parallel decrease in tolerance and *mt1* expression levels in the caeca, but not in the gills. Indeed, Cd toxicokinetics models in *G. fossarum* showed that gills store the metal, while caeca are likely involved in detoxification due to their accumulation and depuration capacities (Gestin et al., 2022). Indeed, it was found that Cd concentrations accumulated in the gills during a 7-day exposure remained stable after a 30-day depuration phase, whereas Cd concentrations in the caeca decreased by 66% (Gestin et al., 2022). As seen previously with the results obtained after field sampling, tolerant gammarids may have acquired, during acclimation processes, a capacity to retain more Cd in the caeca where it can be sequestered by MTs and then eliminated. Our results suggest that Cd tolerance is possibly due to a physiological system in which *mt1* expression in the gills decrease the body-burden of Cd accumulation and the *mt1* expression in the caeca allowed metal detoxification (Figure 3). Moreover, one of the hypotheses that could explain the overexpression of *mt1* in the gills of deacclimatised gammarids from contaminated sites may be that MTs have a shorter half-life than Cd and, as a result, epithelial cells must maintain high levels of transcriptional activity of the *mt1* gene. For example, it has been shown that in the winkle *Littorina littorea*, the half-life of MTs in the gills is 69 days, whereas the half-life of Cd is 300 days (Bebianno and Langston, 1998). The authors of the study therefore suggested a closed metal cycle in the organ, requiring a constant renewal of MTs to sequester the free Cd present in the organ (Bebianno and Langston, 1998). The same observation has also been made in the mussel *Mytilus edulis*, where the half-life of MTs is 25 days compared with 300 days for Cd (Bebianno and Langston, 1993). However, during this same experiment, population B did not show tolerance to Cd (Supplementary Figure 3). This result can be explained by the temporal variability of Cd contamination in this site (Lalouette et al., 2023). Despite this, in this population, high levels of *mt1* expression were found in the gills and caeca, followed by the same dynamics as in the two tolerant populations (A and C population), with a decrease in *mt1* expression in the caeca. Therefore, these results underlined the difficulty to decoupling the exposure parameter from the tolerance parameter and raises questions about temporality where the loss of tolerance appears to be observed more quickly that observed at the molecular level.

We tested the hypothesis that *mt1* expression in offspring depends on parental exposure with the aim of improving our understanding of non-genetic intergenerational transmission of tolerance (*i*.*e*. parental effect) previously described in studies by Vigneron et al. (2019) and Lalouette et al. (2023). For that, Cd tolerance and *mt1* expression were followed in a cohort of neonates born in Cd free water in the laboratory after one reproduction cycle: these neonates resulted from the fertilization of gametes that had formed in the field, but they developed as embryos in the maternal marsupia in Cd-free laboratory conditions. Firstly, Cd tolerance was found in neonates of the three tolerant populations. Interestingly, no difference was observed in the basal level of expression of *mt1* under control conditions between tolerant and sensitive neonates. However, a higher inducibility of *mt1* expression was observed in tolerant neonates after a short re-exposure to Cd, suggesting that is the ability to overexpress *mt1* in contaminated conditions and not constitutive basal levels of *mt1* expression that are responsible for Cd tolerance in *G. fossarum* neonates. In accordance with these observations, Shaw et al. (2019) showed similar results in daphnia (*Daphnia pulex*) after multigenerational exposure to Cd over 50 generations. In this study, identical levels of basal MT expression were found between tolerant and naive individuals after fifty generations and higher MT expression was observed in tolerant individuals after a short re-exposure to Cd (Shaw et al., 2019). Similarly in the worm *Dendrobaena octaedra*, Fisker et al. (2013) showed that the Cd-tolerant individuals of the second-generation exhibited a faster response capacity in the overexpression of a MT during Cd exposure. However, the potential role of MT in the transfer of metal tolerance from adults to offspring has rarely been highlighted, particularly in arthropods. In the study by Guan and Wang (2006), multigenerational exposure to Cd in *Daphnia magna* resulted in an increase in tolerance to this metal and a significant induction of MT in exposed daphnia in all generations, with maximum synthesis in the later generations. The authors then hypothesised that the transgenerational effect of Cd exposure on the rapid induction of MT was due to the increase in Cd concentration in the offspring. Thus, daphnia born from exposed mothers would be more inclined to synthesise MTs (Guan and Wang, 2006). In this same species, another study suggests the role of MTs in the mercury tolerance observed in the offspring since they assume that mRNA for MTs could be transferred through reproduction (Tsui and Wang, 2005).

Finally, in the present study, a symmetrical role was found for both sexes in the transmission of Cd tolerance, correlated with a better capacity for inducing *mt1*. These results support the hypothesis of a link between epigenetics and the transgenerational effect of adaptation to stress in populations (here, the plasticity of Cd sensitivity), as already suggested in our last study (Lalouette et al., 2023) and other authors (Ho and Burggren, 2010; Uller, 2008; Feiner et al., 2022).

## 5. Conclusion

This study conducted in eleven field populations of *G. fossarum* suggests a correlation between the expression of the metallothionein gene, *G. fossarum mt1*, and Cd tolerance plasticity. We have shown that *mt1* gene is over-expressed in the gills and caeca (two organs involved in Cd regulation in this species) in field-tolerant populations. This covariation was also confirmed by a parallel decrease in tolerance and MT expression in the caeca during maintenance in Cd free water in the laboratory. Our study also showed that *mt1* expression regulation can be modulated by historical exposure in field populations and support adaptive processes involving transgenerational mechanisms. These results suggested that links can be drawn between the plasticity of metal detoxification mechanisms at the molecular level (*mt1* expression) and the adaptive plasticity of organism sensitivity at the population level. We suggest that epigenetic mechanisms should be further investigated to improve our understanding of intergeneration acclimation observed in these populations.

## 6. CRediT authorship contribution statement

**Auréline Lalouette:** Writing – original draft, Writing – review & editing, Validation, Methodology, Investigation, Formal analysis, Visualization, Conceptualization. **Arnaud Chaumot:** Writing – review & editing, Validation, Methodology, Funding acquisition, Supervision, Conceptualization. **Louveline Lepeule**: Investigation, Formal analysis. **Karen Gaget** : Resources. **Nicolas Delorme**: Investigation. **Laura Garnero**: Investigation. **Federica Calevro**: Resources. **Davide Degli Esposti**: Writing – review & editing, Validation, Methodology, Supervision, Conceptualization.

## 8. Data Accessibility and Benefit Sharing statement

Raw data on gene expression and acute toxicity will be available upon acceptance in the National data depository https://entrepot.recherche.data.gouv.fr/dataverse/riverly.

## 9. Acknowledgments

The authors benefitted from the French GDR “Aquatic Ecotoxicology” framework which aims at fostering stimulating scientific discussions and collaborations for integrative approaches. The authors thank the ANR (Agence Nationale de la Recherche) program Dyn-Microbiome (ANR-20-CE34-0012). The authors thank Clément Colomb for his assistance in the field experiments especially for the organisms sampling, and Maxence Ruby for his assistance in the molecular experiments.

## Bibliography

Alric, B., Geffard, O., Chandesris, A., Ferréol, M., François, A., Perceval, O., Piffady, J., Villeneuve, B., Chaumot, A., 2019. Multisubstance Indicators Based on Caged Gammarus Bioaccumulation Reveal the Influence of Chemical Contamination on Stream Macroinvertebrate Abundances across France. Environ. Sci. Technol. 53, 5906–5915. 10.1021/acs.est.9b01271

Amiard, J., Amiard-Triquet, C., Barka, S., Pellerin, J., Rainbow, P., 2006. Metallothioneins in aquatic invertebrates: Their role in metal detoxification and their use as biomarkers. Aquatic Toxicology 76, 160–202. 10.1016/j.aquatox.2005.08.015

Amiard-Triquet, C., 2019. Pollution Tolerance in Aquatic Animals: From Fundamental Biological Mechanisms to Ecological Consequences, in: Ecotoxicology. Elsevier, pp. 33–91. 10.1016/B978-1-78548-314-1.50002-X

Atrian, S., Capdevila, M., 2013. Metallothionein-protein interactions. BioMolecular Concepts 4, 143–160. 10.1515/bmc-2012-0049

Bebianno, M.J., Langston, W.J., 1998. Cadmium and metallothionein turnover in different tissues of the gastropod Littorina littorea. Talanta 46, 301–313. 10.1016/S0039-9140(97)00344-5

Bebianno, M.J., Langston, W.J., 1993. Turnover rate of metallothionein and cadmium in Mytilus edulis. Biometals 6, 239–244. 10.1007/BF00187762

Bonneris, E., Giguère, A., Perceval, O., Buronfosse, T., Masson, S., Hare, L., Campbell, P.G.C., 2005. Sub-cellular partitioning of metals (Cd, Cu, Zn) in the gills of a freshwater bivalve, Pyganodon grandis: role of calcium concretions in metal sequestration. Aquatic Toxicology 71, 319–334. 10.1016/j.aquatox.2004.11.025

Chiodi Boudet, L., Mendieta, J., Romero, M.B., Dolagaratz Carricavur, A., Polizzi, P., Marcovecchio, J.E., Gerpe, M., 2019. Strategies for cadmium detoxification in the white shrimp Palaemon argentinus from clean and polluted field locations. Chemosphere 236, 124224. 10.1016/j.chemosphere.2019.06.194

Ciliberti, A., Chaumot, A., Recoura-Massaquant, R., Chandesris, A., François, A., Coquery, M., Ferréol, M., Geffard, O., 2017. Caged Gammarus as biomonitors identifying thresholds of toxic metal bioavailability that affect gammarid densities at the French national scale. Water Research 118, 131– 140. 10.1016/j.watres.2017.04.031

Correia, A.D., Livingstone, D.R., Costa, M.H., 2002. Effects of water-borne copper on metallothionein and lipid peroxidation in the marine amphipod Gammarus locusta. Marine Environmental Research 54, 357–360. 10.1016/S0141-1136(02)00114-9

Dallinger, R., Höckner, M., 2013. Evolutionary concepts in ecotoxicology: tracing the genetic background of differential cadmium sensitivities in invertebrate lineages. Ecotoxicology 22, 767–778. 10.1007/s10646-013-1071-z

Degli Esposti, D., Lalouette, A., Gaget, K., Lepeule, L., Chaabi, Z., Leprêtre, M., Espeyte, A., Delorme, N., Quéau, H., Garnero, L., Calevro, F., Chaumot, A., Geffard., O, 2024. Identification and organ-specific patterns of expression of two metallothioneins in the sentinel species Gammarus fossarum. Comparative Biochemistry and Physiology Part B: Biochemistry and Molecular Biology 110907. 10.1016/j.cbpb.2023.110907

Dong, D.T., Miranda, A.F., Carve, M., Shen, H., Trestrail, C., Dinh, K.V., Nugegoda, D., 2020. Population- and sex-specific sensitivity of the marine amphipod Allorchestes compressa to metal exposure. Ecotoxicology and Environmental Safety 206, 111130. 10.1016/j.ecoenv.2020.111130

Duarte, L.F. de A., Moreno, J.B., Catharino, M.G.M., Moreira, E.G., Trombini, C., Pereira, C.D.S., 2019. Mangrove metal pollution induces biological tolerance to Cd on a crab sentinel species subpopulation. Science of The Total Environment 687, 768–779. 10.1016/j.scitotenv.2019.06.039

Feiner, N., Radersma, R., Vasquez L., Ringnér, M., Nystedt B., Raine A., Tobi E.W., Heijmans B.T., Uller, T., 2022. Environmentally induced DNA methylation is inherited across generations in an aquatic keystone species. iScience 25, 104303. 10.1016/j.isci.2022.104303

Fisker, K.V., Holmstrup, M., Sørensen, J.G., 2013. Variation in metallothionein gene expression is associated with adaptation to copper in the earthworm Dendrobaena octaedra. Comparative Biochemistry and Physiology Part C: Toxicology & Pharmacology 157, 220–226. 10.1016/j.cbpc.2012.11.007

Garret, R.G., 2000. Natural sources of metals to the environment. Human and Ecological Risk Assessment: An International Journal, 6(6), 945–963. 10.1080/10807030091124383

Gestin, O., Lacoue-Labarthe, T., Coquery, M., Delorme, N., Garnero, L., Dherret, L., Ciccia, T., Geffard, O., Lopes, C., 2021. One and multi-compartments toxico-kinetic modeling to understand metals’ organotropism and fate in Gammarus fossarum. Environment International 156, 106625. 10.1016/j.envint.2021.106625

Gestin, O., Lopes, C., Delorme, N., Garnero, L., Geffard, O., Lacoue-Labarthe, T., 2022. Organ-specific accumulation of cadmium and zinc in Gammarus fossarum exposed to environmentally relevant metal concentrations. Environmental Pollution 308, 119625. 10.1016/j.envpol.2022.119625

Gouveia, D., Bonneton, F., Almunia, C., Armengaud, J., Quéau, H., Degli-Esposti, D., Geffard, O., Chaumot, A., 2018. Identification, expression, and endocrine-disruption of three ecdysone-responsive genes in the sentinel species Gammarus fossarum. Science Reports. 8, 3793. 10.1038/s41598-018-22235-7

Guan, R., Wang, W.-X., 2006. Multigenerational cadmium acclimation and biokinetics in Daphnia magna. Environmental Pollution 141, 343–352. 10.1016/j.envpol.2005.08.036

Ho, D.H., Burggren, W.W., 2010. Epigenetics and transgenerational transfer: a physiological perspective. Journal of Experimental Biology 213, 3–16. 10.1242/jeb.019752

Janssens, T.K.S., Roelofs, D., van Straalen, N.M., 2009. Molecular mechanisms of heavy metal tolerance and evolution in invertebrates. Insect Science 16, 3–18. 10.1111/j.1744-7917.2009.00249.x

Khan, F.R., Irving, J.R., Bury, N.R., Hogstrand, C., 2011. Differential tolerance of two Gammarus pulex populations transplanted from different metallogenic regions to a polymetal gradient. Aquatic Toxicology 102, 95–103. 10.1016/j.aquatox.2011.01.001

Lalouette, A., Degli Esposti, D., Garnero, L., Allibert, M., Dherret, L., Dabrin, A., Delorme, N., Recoura-Massaquant, R., Chaumot, A., 2023. Acclimation and transgenerational plasticity support increased cadmium tolerance in Gammarus populations exposed to natural metal contamination in headwater streams. Science of The Total Environment 903. 10.1016/j.scitotenv.2023.166216

Lalouette, A., Degli Esposti, D., Colomb, C., Garnero, L., Quéau, H., Recoura-Massaquant, R., Chaumot, A., 2024. Chronic natural metal contamination shapes the size structure of Gammarus fossarum populations in French headwater rivers. Ecotoxicology. 10.1007/s10646-024-02777-5

Li, S., Sheng, L., Xu, J., Tong, H., Jiang, H., 2016. The induction of metallothioneins during pulsed cadmium exposure to Daphnia magna: Recovery and trans-generational effect. Ecotoxicology and Environmental Safety 126, 71–77. 10.1016/j.ecoenv.2015.10.015

Mao, H., Wang, D.-H., Yang, W.-X., 2012. The involvement of metallothionein in the development of aquatic invertebrate. Aquatic Toxicology 110–111, 208–213. 10.1016/j.aquatox.2012.01.018

Martinez, M., Ramo, J.D., Torreblanca, A., Pastor, A., Diaz-Mayans, J., 1996. Cadmium toxicity, accumulation and metallothionein induction in Echinogammarus echinosetosus. Journal of Environmental Science and Health. Part A: Environmental Science and Engineering and Toxicology 31, 1605–1617. 10.1080/10934529609376445

Mireji, P.O., Keating, J., Hassanali, A., Impoinvil, D.E., Mbogo, C.M., Muturi, M.N., Nyambaka, H., Kenya, E.U., Githure, J.I., Beier, J.C., 2010. Expression of metallothionein and α-tubulin in heavy metal-tolerant Anopheles gambiae sensu stricto (Diptera: Culicidae). Ecotoxicology and Environmental Safety 73, 46–50. 10.1016/j.ecoenv.2009.08.004

Morgan, A.J., Kille, P., Stürzenbaum, S.R., 2007. Microevolution and Ecotoxicology of Metals in Invertebrates. Environ. Sci. Technol. 41, 1085–1096. 10.1021/es061992x

Pedrosa, J., Gravato, C., Campos, D., Cardoso, P., Figueira, E., Nowak, C., Soares, A.M.V.M., Barata, C., Pestana, J.L.T., 2017. Investigating heritability of cadmium tolerance in Chironomus riparius natural populations: A physiological approach. Chemosphere 170, 83–94. 10.1016/j.chemosphere.2016.12.008

Roelofs, D., Janssens, T.K.S., Timmermans, M.J.T.N., Nota, B., Mariën, J., Bochdanovits, Z., Ylstra, B., Van Straalen, N.M., 2009. Adaptive differences in gene expression associated with heavy metal tolerance in the soil arthropod Orchesella cincta. Molecular Ecology 18, 3227–3239. 10.1111/j.1365-294X.2009.04261.x

Rosabal, M., Hare, L., Campbell, P.G.C., 2012. Subcellular metal partitioning in larvae of the insect Chaoborus collected along an environmental metal exposure gradient (Cd, Cu, Ni and Zn). Aquatic Toxicology 120–121, 67–78. 10.1016/j.aquatox.2012.05.001

Shaw, J.R., Colbourne, J.K., Glaholt, S.P., Turner, E., Folt, C.L., Chen, C.Y., 2019. Dynamics of Cadmium Acclimation in Daphnia pulexJ: Linking Fitness Costs, Cross-Tolerance, and Hyper-Induction of Metallothionein. Environ. Sci. Technol. 53, 14670–14678. 10.1021/acs.est.9b05006

Sterenborg, I., Roelofs, D., 2003. Field-selected cadmium tolerance in the springtail Orchesella cincta is correlated with increased metallothionein mRNA expression. Insect Biochem Mol Biol 33, 741–747. 10.1016/s0965-1748(03)00070-5

Stohs, S., 1995. Oxidative mechanisms in the toxicity of metal ions. Free Radical Biology and Medicine 18, 321–336. 10.1016/0891-5849(94)00159-H

Stuhlbacher, A., Maltby, L., 1992. Cadmium resistance in Gammarus pulex (L.). Arch. Environ. Contam. Toxicol. 22. 10.1007/BF00212093

Timmermans, M.J.T.N., Ellers, J., Roelofs, D., van Straalen, N.M., 2005. Metallothionein mRNA Expression and Cadmium Tolerance in Metal-stressed and Reference Populations of the Springtail Orchesella cincta. Ecotoxicology 14, 727–739. 10.1007/s10646-005-0020-x

Tsui, M.T.K., Wang, W.-X., 2005. Influences of maternal exposure on the tolerance and physiological performance of Daphnia magna under mercury stress. Environ Toxicol Chem 24, 1228. 10.1897/04-190R.1

Uller, T., 2008. Developmental plasticity and the evolution of parental effects. Trends in Ecology & Evolution 23, 432–438. 10.1016/j.tree.2008.04.005

Vigneron, A., Geffard, O., Coquery, M., François, A., Quéau, H., Chaumot, A., 2015. Evolution of cadmium tolerance and associated costs in a Gammarus fossarum population inhabiting a low-level contaminated stream. Ecotoxicology 24, 1239–1249. 10.1007/s10646-015-1491-z

Vigneron, A., Geffard, O., Quéau, H., François, A., Chaumot, A., 2019. Nongenetic inheritance of increased Cd tolerance in a field Gammarus fossarum population: Parental exposure steers offspring sensitivity. Aquatic Toxicology 209, 91–98. 10.1016/j.aquatox.2019.02.001

Vijver, M.G., van Gestel, C.A.M., Lanno, R.P., van Straalen, N.M., Peijnenburg, W.J.G.M., 2004. Internal Metal Sequestration and Its Ecotoxicological Relevance: A Review. Environ. Sci. Technol. 38, 4705–4712. 10.1021/es040354g

Weng, N., Wang, W.-X., 2014. Improved tolerance of metals in contaminated oyster larvae. Aquatic Toxicology 146, 61–69. 10.1016/j.aquatox.2013.10.036

Wu, S.M., Lin, H., Yang, W., 2008. The effects of maternal Cd on the metallothionein expression in tilapia (Oreochromis mossambicus) embryos and larvae. Aquatic Toxicology 87, 296–302. 10.1016/j.aquatox.2008.02.012

Wu, S.M., Tsai, P.R., Yan, C.J., 2012. Maternal cadmium exposure induces mt2 and smtB mRNA expression in zebrafish (Danio rerio) females and their offspring. Comparative Biochemistry and Physiology Part C: Toxicology & Pharmacology 156, 1–6. 10.1016/j.cbpc.2012.02.001

Ziller, A., Fraissinet-Tachet, L., 2018. Metallothionein diversity and distribution in the tree of life: a multifunctional protein. Metallomics 10. 10.1039/C8MT00165K

