## Supplementary data for "Cadmium tolerance is associated with tissue-specific plasticity of metallothionein gene expression in *Gammarus fossarum* field populations"

### Supplementary material

#### Supplementary Tables:

**Supplementary Table 1 :** Localisation and characteristics of the 11 sites. The status towards bioavailable Cd contamination was defined following an active biomonitoring approach based on *Gammarus* caging. Here, we present the average bioaccumulated Cd concentrations during different tests conducted over 2015-2022 period, data already reported in Lalouette et al. (2024). Concentrations of metals and bioavailable background assessment concentrations (BBAC) are expressed in µg/g dw (dry weight). BBAC corresponds to the 95 % percentile of the national distribution of the background concentrations recorded in French rivers (Alric et al., 2019). According to Lalouette et al. (2023), the status of population Cd-exposure was defined as followed: "contaminated" if accumulated concentration is higher than BBAC for Cd (0.3 µg Cd/g), "intermediate" if the Cd level is between 0.2 and 0.3 µg Cd/g, and "uncontaminated" if the level is less than 0.2 µg Cd/g.

| Site | Name of the river | Location (city, district, country) | GPS coordinates | Characterisation of the bioavailable Cd contamination | |
|---|---|---|---|---|---|
| | | | | Mean bioavailable Cd contamination<br>*BBAC = 0.30* | Status towards bioavailable Cd contamination |
| Ardillats (A) | L'Ardières | Les Ardillats, Rhône, France | 46°11'12.0"N 4°31'16.0"E | 0.66 | Contaminated |
| Marchampt (B) | Marchampt | Marchampt, Rhône, France | 46°07'09.0"N 4°32'52.0"E | 0.39 | Contaminated |
| Vernay (C) | Vernay | Vernay, Rhône, France | 46°10'07.0"N 4°31'44.0"E | 0.52 | Contaminated |
| Strengbach (D) | Strengbach | Aubure, Haut-Rhin, France | 48°12'37.8"N 7°12'55.4"E | 0.29 | Intermediate |
| Rauental (E) | Rauenthal | Sainte-Marie-aux-Mines, Bas-Rhin, France | 48°13'40.4"N 7°09'57.2"E | 0.24 | Intermediate |
| Morcille (F) | La Morcille | Villié-Morgon, Rhône, France | 46°10'42.2"N 4°38'06.7"E | 0.22 | Intermediate |
| Seran (G) | Le Séran | Béon, Ain, France | 45°51'21.2"N 5°43'14.5"E | 0.08 | Uncontaminated |
| Doulonne (H) | La Doulonne | Plumont, Jura, France | 47°07'13.8"N 5°44'07.8"E | 0.16 | Uncontaminated |
| Katlen (I) | Katlen | Lièpvre, Haut-Rhin, France | 48°16'20.6"N 7°20'20.8"E | 0.09 | Uncontaminated |
| Reigne (J) | La Reigne | Magny-Vernois, Haute-Saône, France | 47°39'23.0"N 6°28'00.1"E | 0.15 | Uncontaminated |
| Vancelle (K) | Vancelle | Vancelle, Bas-Rhin, France | 48°16'52.0"N 7°18'04.7"E | 0.07 | Uncontaminated |

**Supplementary Table 2 :** Identification of the metallothionein MT-1 in the amphipod *Gammarus fossarum* (modify from Degli-Esposti et al. 2024)

| IDs | Coding DNA Sequence | Primers qPCR |
|---|---|---|
| GFBF_DN110944_c4_g1_i2<br>putative MT-1 | ATGCCTAACGACTGCTGCAAAGAGGACTGC<br>AAATGCACCGCCGAGGAATGCGGGAAGGAC<br>TGCGGCTGCACGGACTGCGACTGCCAGAAG<br>TGCGAGAATTGCAAGGGCAGCTGCGACTGCT<br>CCTCTGTTGACGCCTGCGCTACCAACTGCGA<br>CACGCCATGCAGCTGCTGCCCTACGGAGTAG | Forward:<br>AACGACTGCTGCAAAGAGGA<br>Reverse:<br>CAGCTGCCCTTGCAATTCTC |

**Supplementary Table 3 :** Statistical overview of the Cd tolerance of male adults from populations A. B. C. G and H after sampling in the field (T0) and after their maintenance during 2 months in Cd-free water (T2). Adults were exposed to 80 µg/L of Cd.

| Population | Number of adults | Median survival time LT50 (days) [95% confidence interval] | 25th percentile of adults survival time (days) | 75th Percentile of adults survival time (days) | % of adults with survival time ≤ 6 days |
|---|---|---|---|---|---|
| A-T0 | 45 | 12 [9 ; 18] | 7 | 17 | 15.5 |
| B-T0 | 45 | 6 [5 ; 9] | 4 | 10 | 55.5 |
| C-T0 | 45 | 18 [13 ; 21] | 9 | 21 | 13.3 |
| G-T0 | 45 | 8 [7 ; 9] | 6 | 9 | 28.8 |
| H-T0 | 45 | 7 [6 ; 10] | 5 | 11 | 37.7 |
| A-T2 | 45 | 8 [6 ; 14] | 5 | 14 | 40 |
| B-T2 | 45 | 7 [6 ; 9] | 5 | 10 | 42.2 |
| C-T2 | 45 | 7 [6 ; 9] | 5 | 10 | 44.4 |
| G-T2 | 45 | 6 [5 ; 7] | 4 | 7 | 62.2 |
| H-T2 | 45 | 8 [7 ; 9] | 6 | 10 | 26.6 |

**Supplementary Table 4 :** Statistical overview of *G. fossarum mt1* fold-change in neonates from the tolerant population (AA). the sensitive population (GG) and from population crossing (AG and GA) after re-exposure at 3 µgCd/L during 3 days.

| Condition (♂/♀) | Number of neonates | Fold-change | Minimum | Maximum | % of neonates with fold-change ≥ 4 |
|---|---|---|---|---|---|
| AA | 6 | 9.2 | 6.9 | 12.1 | 100 |
| AG | 6 | 5.0 | 2.6 | 6.7 | 67 |
| GA | 6 | 3.7 | 2.1 | 5.9 | 50 |
| GG | 6 | 3.7 | 2.6 | 4.2 | 17 |

#### Supplementary Figures:

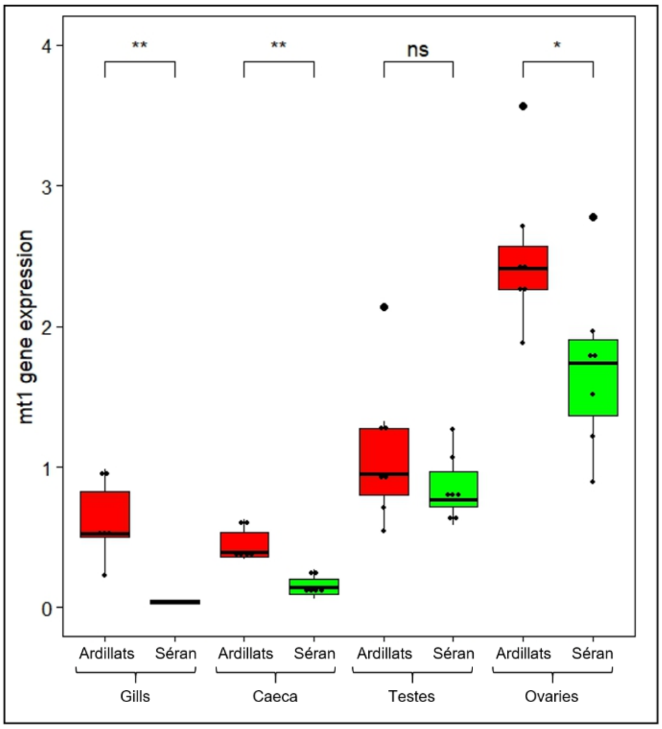


**Supplementary Figure 1 :** *G. fossarum mt1* gene expression measured by RT-qPCR (√ΔCt) in the gills. caeca. testes and ovaries of adult after sampled in the field in Ardillats and Séran population. Red and green colors correspond respectively to Cd contaminated and uncontaminated status of populations. Gene expression was normalize using the elongation factor (EF) Ct value. The symbols indicate the statistical significance (p-value) as followed: (ns) > 0.05; (*) ≤ 0.05; (**) ≤ 0.01; (***) ≤ 0.001; (****) ≤ 0.0001.

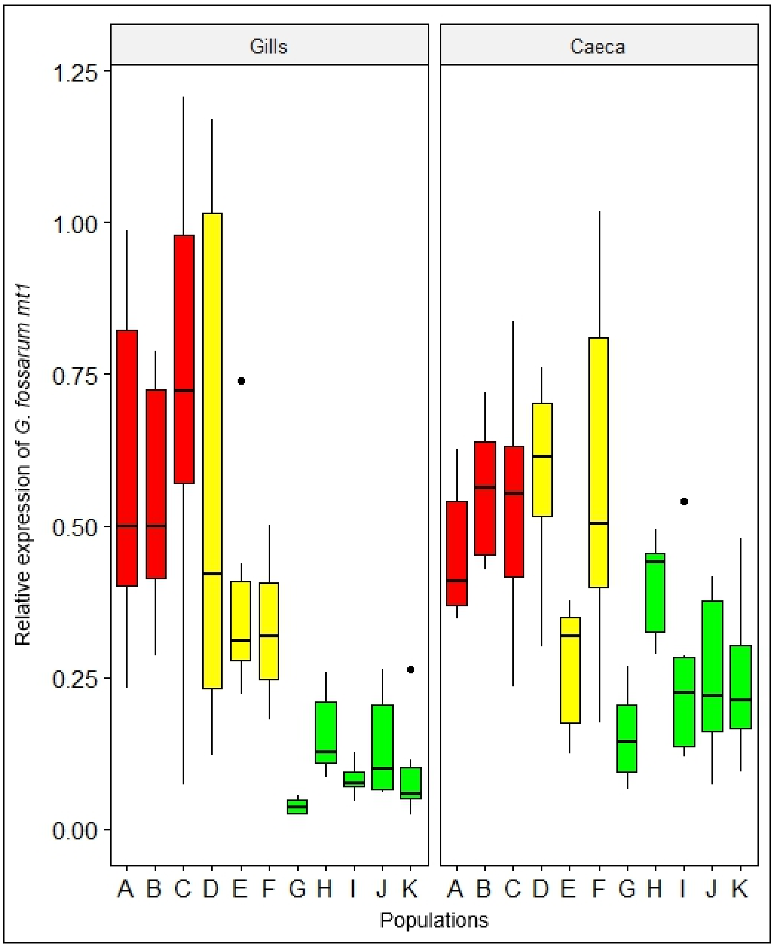


**Supplementary Figure 2 :** *G. fossarum mt1* gene expression measured by RT-qPCR (√ΔCt) in the gills and caeca of adult males after sampled in the field in the 11 populations (A. B. C. D. E. F. G. H. I. J. K). Red. yellow and green colors correspond respectively to Cd contaminated. intermediate and uncontaminated status of populations.

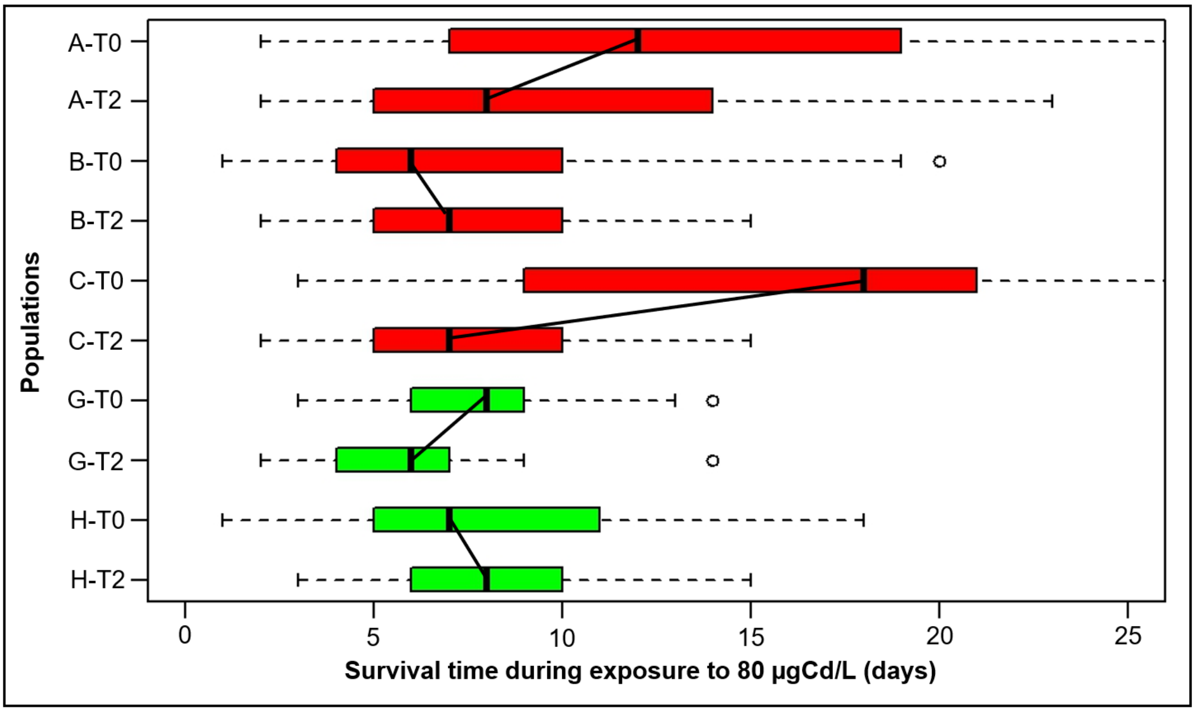


**Supplementary Figure 3** : Evolution of Cd tolerance of field populations (A. B. C. G. H) during maintenance in Cd-free conditions. The boxplots present the distribution of individual survival times of male adults after sampling in the field (T0) and after their maintenance during 2 months in Cd-free water (T2). Adults were exposed to 80 µg/L of Cd. Red and green colors correspond respectively to Cd contaminated and uncontaminated status of populations in their habitat of origin.

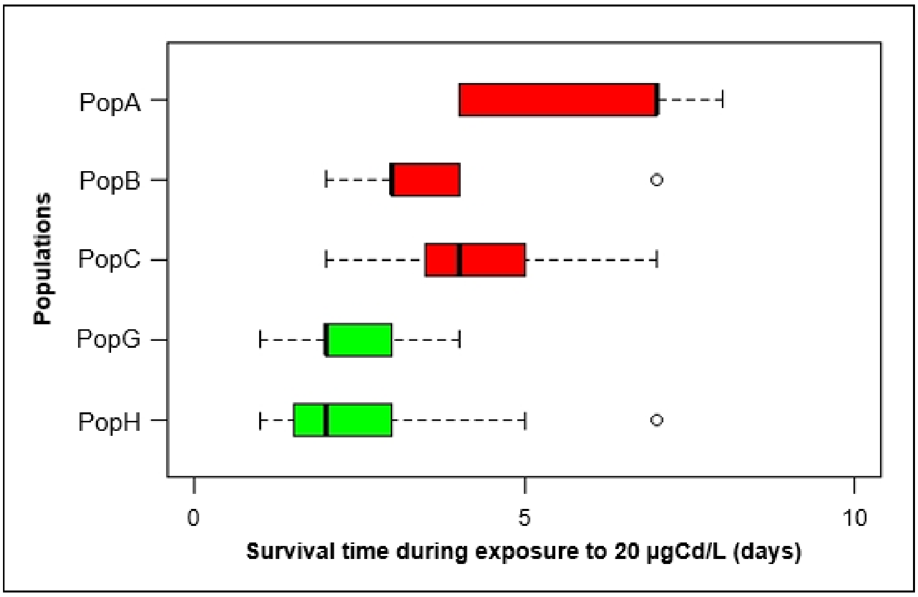


**Supplementary Figure 4** : Cd tolerance in the offspring of five populations (A. B. C. G. H) born in the lab in Cd free water after one reproduction cycle (approximatively 25 days). Neonates were exposed to Cd individually one day after their release from maternal marsupium. Red and green colors correspond respectively to Cd contaminated *vs* uncontaminated status of populations in their habitat of origin.

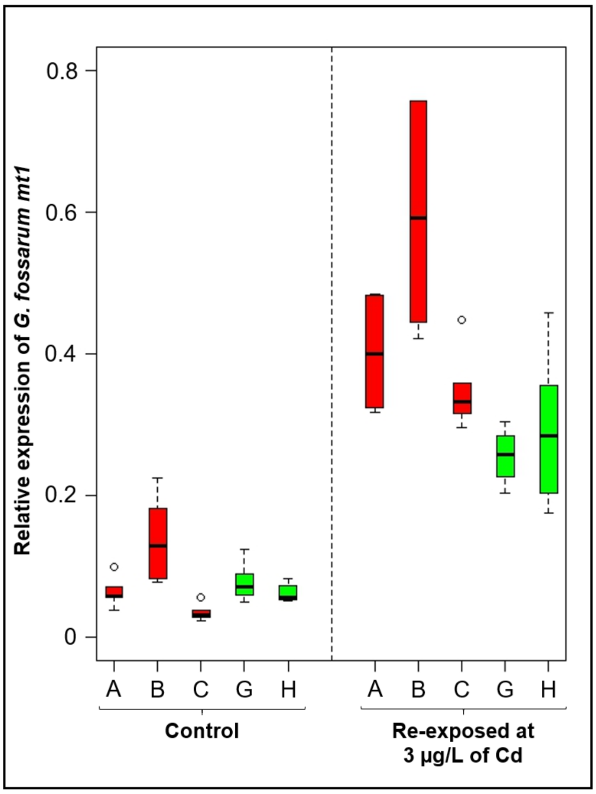


**Supplementary Figure 5** : *G. fossarum mt1* gene expression measured by RT-qPCR (√ΔCt) in neonates born in the lab in Cd free water after one reproduction cycle (approximatively 25 days) without Cd exposure and after a re-exposure at 3 µgCd/L during 3 days. Red and green colors correspond respectively to Cd contaminated and uncontaminated status of populations in their habitat of origin.

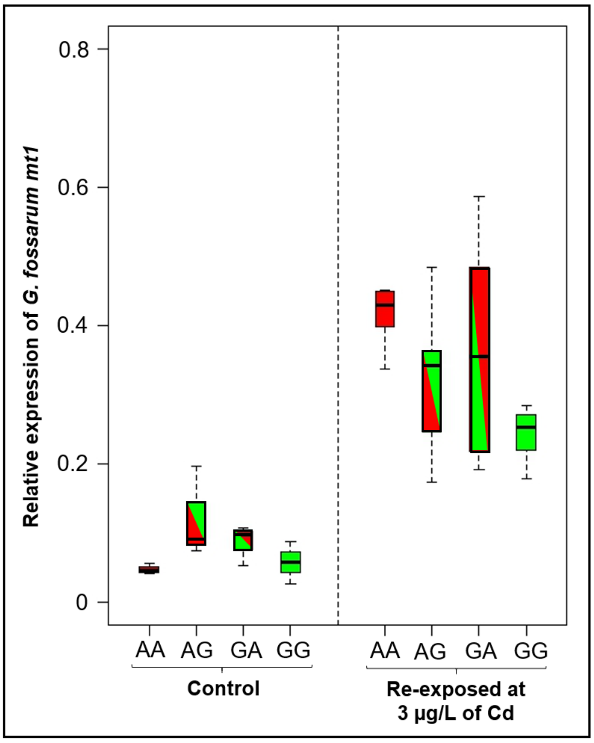


**Supplementary Figure 6** : *G. fossarum mt1* gene expression measured by RT-qPCR (√ΔCt) in neonates from the tolerant population (AA). the sensitive population (GG) and from population crossing (AG and GA) without Cd exposure and after re-exposure at 3 µgCd/L during 3 days
